# Frequency-Dependent Modulation of Adult Hippocampal Neurogenesis, Memory, and BDNF Signaling by Low-Intensity Focused Ultrasound

**DOI:** 10.64898/2026.05.09.723959

**Authors:** Kareen Kanaan, Heba Badawe, Wassim Abu-Khier, Massoud Khraiche

## Abstract

Adult hippocampal neurogenesis plays a central role in learning, memory formation, and adaptive neural plasticity, making it an attractive target for noninvasive neuromodulation strategies. Low-intensity focused ultrasound (LIFU) has emerged as a promising modality for modulating brain function, yet its effects on adult neurogenesis and the role of stimulation frequency remain incompletely understood. In this study, we evaluated whether transcranial LIFU applied to the dentate gyrus influences neurogenic and cognitive outcomes in a frequency-dependent manner. Adult rats received twice-weekly ultrasound stimulation for four weeks at 0.5, 1, or 5 MHz. Neurogenesis was assessed through BrdU incorporation and neuronal differentiation by BrdU/NeuN co-labeling, while expression of neurogenesis-associated markers (BDNF, FGF-2, and Sox-2) was quantified using qRT-PCR. Behavioral effects were examined using the novel object recognition task. Among the tested conditions, 0.5 MHz stimulation produced the most pronounced neurogenic response, with increased cellular proliferation in the dentate gyrus, elevated expression of neurogenic markers, and improved recognition memory relative to sham-treated animals. Higher stimulation frequencies yielded comparatively weaker effects. These findings identify stimulation frequency as a critical determinant of LIFU-driven neuroplastic responses and support the potential of focused ultrasound as a noninvasive approach for promoting hippocampal regeneration and functional recovery.

## Introduction

Neurodegenerative disorders such as Alzheimer’s disease (AD) and Parkinson’s disease (PD) are progressive conditions characterized by neuronal loss and dysfunction, resulting in cognitive decline, motor impairment, and behavioral disturbances [1]. A key factor contributing to their onset and progression is the age-related decline in the brain’s capacity for neuronal regeneration and axonal repair, particularly the reduction of adult hippocampal neurogenesis [2-4]. In the adult mammalian brain, neurogenesis is largely confined to the subgranular zone (SGZ) of the hippocampal dentate gyrus and the subventricular zone (SVZ) of the lateral ventricles. Within these neurogenic niches, neural progenitors generate new neurons that integrate into existing circuits to support memory, learning, and behavioral flexibility [5]. However, no approved therapy currently promotes neural regeneration or restores cognitive function in affected individuals. This therapeutic gap underlines the urgent need for novel regenerative strategies.

Low-intensity focused ultrasound (LIFU) is a non-invasive neuromodulatory modality capable of targeting deep brain structures with millimeter precision and minimal tissue damage [6-8]. Preclinical studies indicate that LIFU can modulate neural activity, induce synaptic plasticity, and stimulate the release of neurotrophic factors such as brain-derived neurotrophic factor (BDNF) and fibroblast growth factor-2 (FGF-2) [9-11]. These factors are central to creating a neurogenic microenvironment, supporting neural stem cell proliferation and survival, and maintaining hippocampal circuit integrity. Importantly, LIFU has also been reported to stimulate neural stem cell proliferation and facilitate blood-brain barrier permeability, highlighting its potential as a regenerative intervention [12]. Indeed, recent studies have linked LIFU stimulation of the hippocampus to enhanced neurogenesis and improved memory performance in rodent models of aging and neurodegeneration, though the precise mechanisms remain under investigation [13-15].

Recent studies have shown that low-intensity ultrasound can modulate neuroinflammation, synaptic activity, neurotrophic signaling, and behavior across multiple CNS contexts [16-20]. Accordingly, the present study was designed as a controlled parametric investigation to isolate the role of ultrasound frequency as a biophysical determinant of adult hippocampal neurogenesis and cognitive function, rather than as a mechanistic dissection of downstream signaling pathways. Despite these advances, the regenerative capacity of LIFU remains insufficiently characterized, particularly with respect to stimulation frequency [21]. Frequency is a critical acoustic parameter: lower frequencies (≤1 MHz) penetrate deeper with less attenuation, potentially delivering stronger mechanical stimulation to subcortical neurogenic regions [22].

Mechanical stimulation has been shown to regulate neural progenitor proliferation through calcium-dependent signaling pathways, including activation of mechanosensitive ion channels such as Piezo1 and TRP family members. Downstream engagement of MAPK/ERK and CREB signaling cascades promotes transcription of neurotrophic factors including BDNF, which are critical for adult hippocampal neurogenesis. Whether ultrasound frequency differentially engages these pathways remains unknown.

In this study, we examined the frequency-dependent effects of transcranial LIFU (0.5, 1, and 5 MHz) on adult hippocampal neurogenesis and cognitive performance in rats. We hypothesized that low-frequency ultrasound would more effectively enhance neural progenitor proliferation and activate neurotrophic pathways compared to higher frequencies. To test this, we applied LIFU to the dentate gyrus twice weekly for four weeks and assessed outcomes using BrdU labeling, mRNA expression of neurogenic regulators (BDNF, FGF-2, Sox-2), and behavioral performance in the novel object recognition (NOR) test. We therefore performed a systematic, within-study comparison of multiple ultrasound frequencies (0.5, 1, and 5 MHz) under matched transcranial exposure conditions to determine whether frequency alone governs neurogenic and behavioral outcomes in vivo. This study aims to identify frequency-dependent biological responses, providing a framework for future mechanistic investigations rather than directly establishing molecular causality.

## Materials and Methods

### Animal Preparation

Adult male Sprague-Dawley rats (12 weeks old, 500–600 g) were used in this study. This strain was selected for its large size, which facilitates surgical precision and experimental handling, as well as its genetic uniformity, which reduces inter-animal variability. The 12-week age corresponds to full brain maturity and stable cognitive traits, making it suitable for investigating adult hippocampal neurogenesis. Male rats were used to avoid hormonal fluctuations associated with the female estrous cycle, which could confound behavioral and neurogenic outcomes. Animals were housed under standard laboratory conditions (12-hour light/dark cycle, 20–22°C), with ad libitum access to food and water. All animals were acclimated for at least one week prior to experimentation. Anesthesia was induced with an intraperitoneal injection of ketamine (80 mg/kg) and xylazine (20 mg/kg) prior to all procedures[23, 24]. Rats were euthanized at the end of the treatment period for molecular and histological analyses. All protocols were approved by the Institutional Animal Care and Use Committee (IACUC) of the American University of Beirut.

### Experimental Design, Stereotaxy and Ultrasound Stimulation Protocol

Group allocation was performed using a computer-generated randomization sequence assigned into five groups: naïve (n = 3), sham (n = 3), and three experimental groups (n = 5 per group) exposed to LIFU at different frequencies (0.5 MHz, 1 MHz, and 5 MHz). The naïve group received no treatment. The sham group underwent anesthesia and stereotaxic positioning without ultrasound exposure to control for handling and anesthesia-related effects. Experimental groups received transcranial LIFU targeting the hippocampus using single-element focused transducers. Specifically, the 0.5 MHz transducer from Mana Instruments (E0525-SU) had a diameter of 25 mm and a focal length of 30 mm. For higher frequencies, Olympus transducers (Olympus NDT, USA), model V314-SU-F (1 MHz) and V308-SU-F (5 MHz), were used; both have a diameter of 19.05 mm and focal lengths of 37.19 mm and 39.52 mm, respectively. Sample sizes were determined based on prior studies examining ultrasound-induced modulation of hippocampal plasticity and were sufficient to detect biologically meaningful differences in BrdU labeling and gene expression.

For precise targeting, anesthetized animals were secured in a stereotaxic frame to ensure stable head fixation. The scalp was shaved to optimize acoustic coupling and transducer placement. To maintain a fully noninvasive protocol, the skull was not surgically exposed. Instead, Bregma was approximated externally using cranial landmarks, aligned with the interaural line. Specifically, bregma was estimated at 9.0 mm anterior and 10.0 ± 0.2 mm dorsal to the interaural zero plane. This external approximation was used as a reference point for stereotaxic positioning of the ultrasound transducer. The dorsal hippocampus was targeted along the rostrocaudal axis from –2.12 mm to –6.3 mm relative to bregma (Fig. 1A) [25]. Once aligned, a layer of ultrasound coupling gel was applied between the transducer face and the shaved scalp to facilitate optimal acoustic transmission. The transducer was then positioned according to the calculated coordinates for hippocampal stimulation. Ultrasound stimulation was delivered at a spatial-peak pulse-average intensity (ISPPA) of 3 W/cm^2^, with a 5% duty cycle and a 20 Hz pulse repetition frequency. Each session lasted 5 minutes and was administered twice weekly for four weeks. Acoustic field simulations were performed using the k-Wave MATLAB toolbox [26-29], which solves acoustic wave propagation in heterogeneous, absorbing media using a k-space pseudo-spectral method. We modeled ultrasound propagation through a 2D medium consisting of a water-skull interface to estimate the acoustic field profiles for each transducer frequency (0.5, 1, and 5 MHz). The computational domain was a 64 x 128 grid with a 0.3 mm step size and a perfectly matched layer (PML) of 20 grid points on all boundaries to suppress reflections [30, 31]. Each transducer was modeled as a spherically focused source with a circular aperture and a focal length matching its physical specifications. The source was driven at its corresponding center frequency using a sinusoidal tone burst. Acoustic properties of the rat skull (density: ρ = 1732 kg/m^3^; sound speed: c = 2850 m/s) were defined homogeneously across a 0.71 mm-thick skull layer, based on literature-reported values and defined in a piecewise manner across the simulation grid [32]. A sensor mask was positioned along the beam axis and focal zone to record the spatiotemporal pressure distribution. Simulations were conducted with and without the skull layer to quantify acoustic attenuation and focal distortion[33]. The resulting intensity profiles were used to calculate the frequency-dependent attenuation coefficient α, based on the logarithmic ratio of maximum intensities in water and skull conditions according to the following equation [34, 35]: 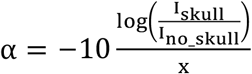, where I_no_skull_ and I_skull_ represent the peak spatial intensities recorded with and without the skull respectively, and x represents the skull thickness of 0.71 mm. Attenuation values are summarized in table 1, and representative beam profiles for each frequency are shown in Fig. 1B. These simulations highlight frequency-dependent beam distortion, attenuation, and focal shifts, underscoring the importance of accounting for skull effects in transcranial ultrasound targeting. Based on the calculated acoustic attenuation at each ultrasound frequency, the output amplitude of the waveform generator was adjusted to ensure that the in-skull spatial peak pulse-average intensity (I_skull_) was maintained at 3 W/cm^2^ in all the trials.

**Table 1.** Ultrasound intensity with and without skull tissue, and corresponding attenuation coefficients (α) across frequencies.

| Frequency | $I_{\text{no\_skull}}$ (W/cm<sup>2</sup>) | $I_{\text{skull}}$ (W/cm<sup>2</sup>) | $\alpha$ (dB/cm) |
| --- | --- | --- | --- |
| 0.5 MHz | 2.367 | 1.949 | 11.81 |
| 1 MHz | 3.117 | 2.4599 | 14.39 |
| 5 MHz | 8.909 | 3.314 | 60.12 |

**Fig. 1.**
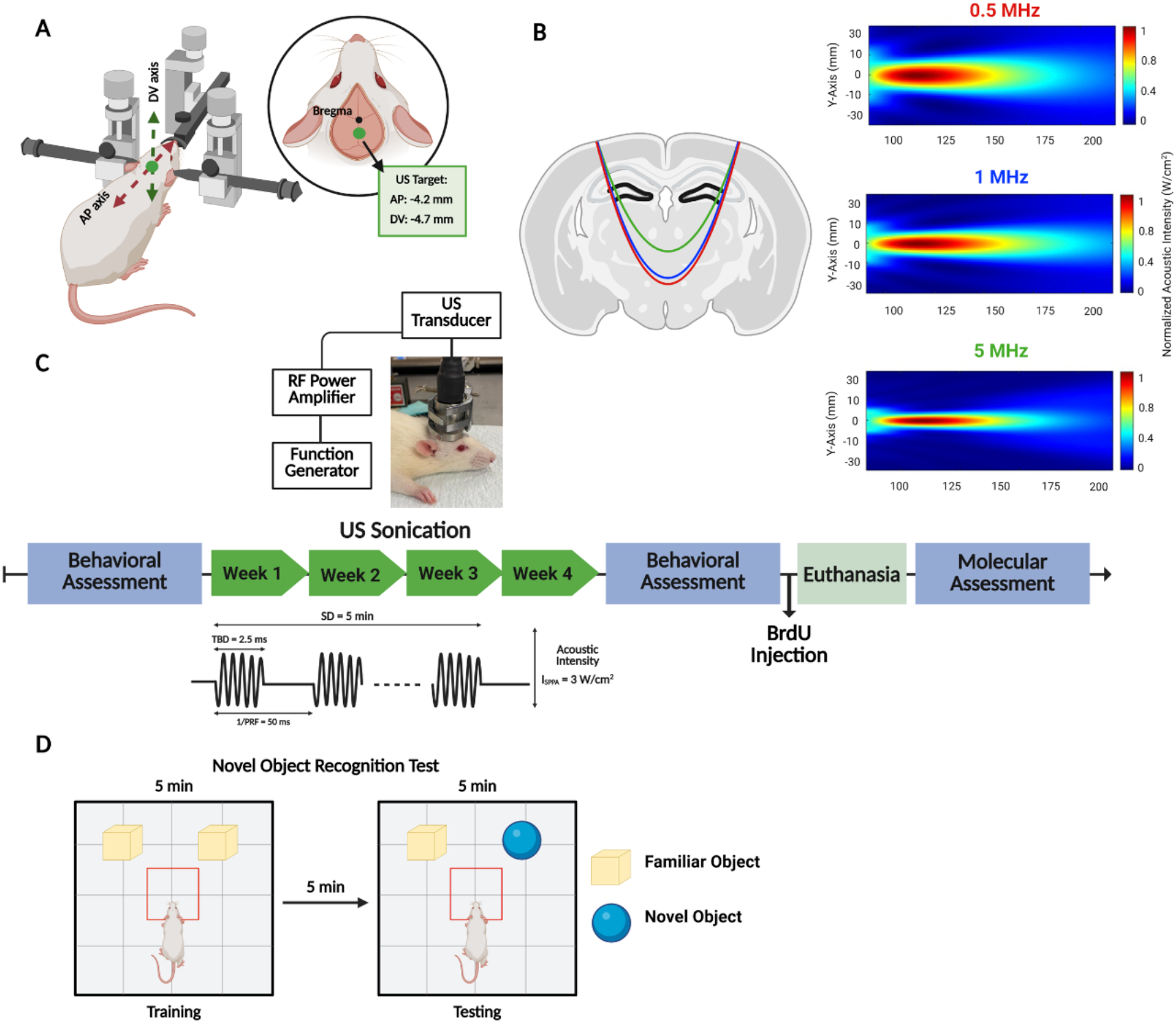
Experimental protocol. (A) FUS was stereotaxically targeted to the dentate gyrus of the hippocampus (coordinates: AP = –4.2 mm, DV = –4.7 mm relative to Bregma). (B) Simulations of the acoustic intensity profile across the longitudinal plane of the three ultrasound transducers used (right). A coronal section of the rat brain marked by three semi-ellipses which present -3dB sonication region of each ultrasound frequency used (left). (C) Experimental timeline where the rats underwent baseline behavioral assessment followed by four weeks of ultrasound stimulation. Each session consisted of 5 min sonication at an acoustic intensity of 3 W/cm^2^ (Isppa), tone-burst duration (TBD) of 2.5 ms, and a pulse repetition frequency (PRF) of 20 Hz. Behavioral assessments were repeated after the stimulation period, followed by BrdU injection, euthanasia, and molecular analyses. (D) Schematic diagram for the novel object recognition test.

### Behavioral Assessment

On Day 0 (Baseline), animals underwent behavioral assessment using the novel object recognition task to evaluate baseline cognitive performance. Following the four-week treatment phase, the NOR task was repeated to assess post-treatment cognitive changes (Fig. 1C). For cellular analysis, all animals received a single intraperitoneal injection of 5-bromo-2’-deoxyuridine (BrdU), a thymidine analog that labels dividing cells during the S-phase of the cell cycle and serves as a marker for cell proliferation. Animals were euthanized 24 hours later, and brain tissues were collected and processed for cellular and molecular analyses.

The NOR task was conducted in the open-field apparatus at both time points (baseline and post-treatment) to assess recognition memory [36, 37]. The test consisted of two phases: a familiarization phase and a testing phase (Fig 1D). During the familiarization phase, two identical objects were placed in opposite corners of the open field. Each rat was placed facing the opposite side of the objects and allowed to explore freely for 5 minutes. After a 5-minute inter-phase interval, during which the rat was returned to its home cage and the field was cleaned with 70% ethanol, the testing phase was initiated. In this phase, one of the familiar objects was replaced with a novel object, while the spatial arrangement remained unchanged. The rat was then reintroduced to the field for additional 5 minutes of exploration. Behavioral performance was recorded and analyzed using AnyMaze^™^ tracking software (Fort Worth, TX, USA). The primary parameters quantified included the time spent in the novel versus familiar object zones, the time spent oriented toward the novel object, and the time getting closer to the novel object. Object recognition memory was assessed by quantifying the animal’s tendency to spend more time exploring novel objects compared to familiar ones.

### Cellular Assessment

Following ultrasound stimulation, cellular proliferation was assessed. BrdU powder (Sigma-Aldrich, B5002) was dissolved in 0.9% warm sterile saline to achieve a final dose of 200 mg/kg. A 900 µL injection was given to each rat. The rats received a total of 3 BrdU injections (66 mg/kg per 300 µL injection, intraperitoneally), with each injection spaced 2 hours apart. For tissue sampling and processing, the rats were deeply anesthetized with intraperitoneal injections of ketamine (Ketalar^®^; 50 mg/kg) and Xylazine (Xylazine^®^; 12 mg/Kg), followed by transcardial perfusion. The brains were extracted and fixed in 4% paraformaldehyde for 24 hours, then cryoprotected in 30% sucrose in 0.1 M phosphate-buffered saline (PBS) at 4 °C until complete impregnation. Brain sections were cut using a sliding microtome and systematically sampled into six sets using the fractionator method [38, 39]. Specifically, 40 µm coronal sections were serially cut from the rostral to caudal extent of the dentate gyrus (DG) at the following rostro-caudal coordinates to cover the entire hippocampal formation: −2.12 to −6.3 mm relative to bregma. To map the BrdU distribution topographically, the DG was divided into three regions: the rostral region from −2.12 to −3.7 mm relative to bregma, the intermediate region from −3.7 to −4.9 mm, and the caudal region from −4.9 to −6.3 mm [25, 39]. The rostral region corresponds to the dorsal hippocampus, while the intermediate and caudal regions are part of the ventral hippocampus. All sections were stored in 15 mM sodium azide solution in 0.1 M PBS until processing.

For immunofluorescence staining, sections were washed and incubated at 37 °C with 2N HCL for DNA denaturation, followed by neutralization with 0.1M sodium borate buffer (pH 8.5) for 10 minutes. Sections were then incubated overnight at 4 °C with monoclonal primary antibodies: mouse anti-BrdU (1:500; Santa Cruz) and rabbit anti-NeuN (1:500; Millipore) diluted in PBS containing 3% normal goat serum (NGS), 3% bovine serum albumin (BSA), and 0.1% Triton-X. Incubation with secondary antibodies would follow on the second day for 2 hours on a shaker at room temperature using Alexa Fluor-568 goat anti-mouse (1:250; Molecular Probes, Invitrogen) and Alexa Fluor-488 goat anti-rabbit (1:250; Molecular Probes, Invitrogen). Finally, sections were washed and mounted onto glass slides with Fluoro-Gel mounting medium containing DAPI (Electron Microscopy Sciences, USA).

BrdU+ cells were counted under a 40× oil immersion objective within the SGZ of the DG to quantify cell proliferation. One random set of sections was selected per animal. The number of BrdU+ cells per section was multiplied by 6 (total number of sets per rat) to estimate the total number of BrdU+ cells per region (rostral, intermediate, and caudal) per animal. Confocal imaging was performed using a Zeiss LSM 710 laser scanning confocal microscope with a 40× oil objective. Z-stacks and tile scans were acquired to visualize the full 40 µm thickness of the DG in each region. Images were processed using Zeiss ZEN 2009 software with maximum intensity projection. All cell quantification and image analyses were performed by an investigator blinded to experimental group allocation. Behavioral scoring and molecular analyses were conducted using coded samples to ensure blinded assessment.

### Molecular Assessment: RNA Extraction and Quantitative Real-Time PCR

For molecular assessment, total RNA was extracted from hippocampal tissues using TriZol (TRI reagent®, Sigma; St. Louis, MO, USA) according to the manufacturer’s instructions. Briefly, tissue was homogenized in 1 mL of TriZol reagent on ice, followed by addition of 0.2 mL chloroform and centrifugation at 12,000 rpm for 20 minutes at 4°C. The RNA-containing phase was mixed with 0.35 mL of isopropanol and incubated at room temperature for 10 minutes, then centrifuged at 15,000 rpm for 30 minutes. The RNA pellet was washed twice with 70% ethanol, air-dried, and then resuspended in RNase-free water. RNA concentration and purity were measured using a ThermoScientific^™^ NanoDrop 2000^™^, and RNA integrity was verified using the Agilent BioAnalyzer 2100™. cDNA synthesis was performed using the QIAGEN QuantiTect reverse transcription kit, according to the manufacturer’s instructions. The resulting cDNA was diluted at a 1:10 ratio for downstream applications. The mRNA expression levels of hippocampal samples from all groups were measured by RT-PCR (Bio-rad CFX^™^ Manager Software; cat #1845000; Hercules, CA, USA) using the ΔΔCt method with SYBR Green system (Applied Biosystems; cat #A46111; Waltham, MA, USA). The PCR protocol included an initial denaturation at 95°C for 5 minutes, followed by 40 cycles of denaturation at 95°C for 10 seconds, annealing at 57°C for 30 seconds, and extension at 72°C for 10 minutes. Each hippocampal sample from an individual animal was considered one biological replicate. qPCR reactions for each biological replicate were run in technical duplicate, and the mean Ct value was used for ΔΔCt analysis. Target gene expression was normalized to the housekeeping gene GAPDH. Relative gene expression was quantified using the following equation:

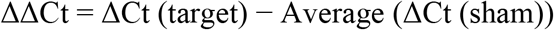

where ΔCt = Ct (target) − Ct (GAPDH). The relative expression of the target gene compared to GAPDH was calculated as 2^−ΔΔCt^. A list of primers and their sequences is provided in Table 2.

**Table 2.**
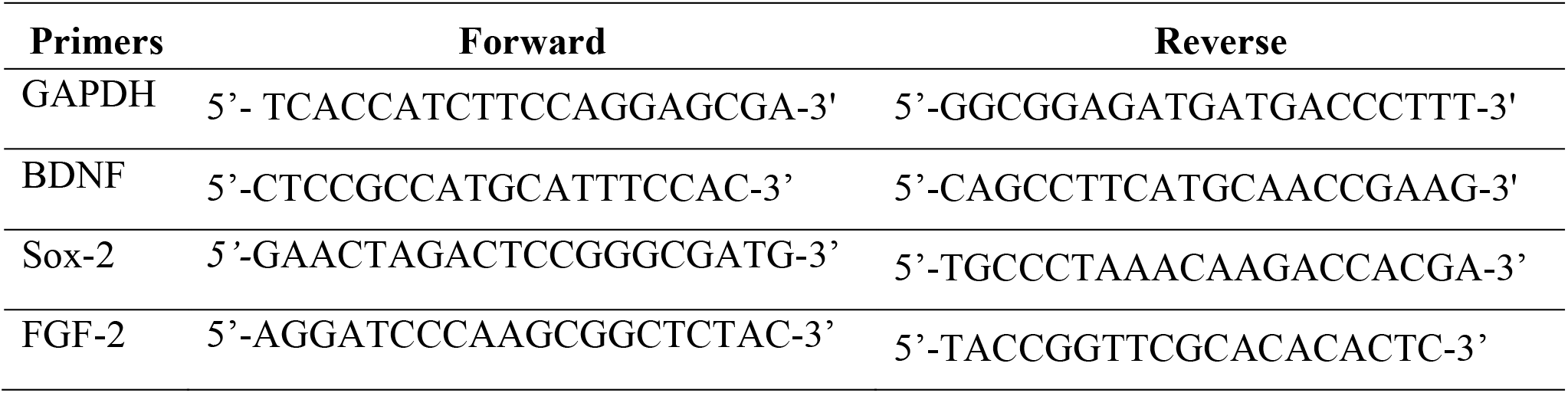
Primers used with qRT-PCR in the study. Brain-derived neurotrophic factor: BDNF; SRY-box transcription factor 2: Sox-2; Fibroblast growth factor 2: FGF-2.

## Statistical Analysis

Statistical analyses were performed using GraphPad Prism 9 (GraphPad Software, Inc., CA, USA). Data distribution was assessed using the Shapiro–Wilk test for normality, and homogeneity of variance was evaluated using Levene’s test. For comparisons among multiple groups, one-way ANOVA was conducted followed by Tukey’s post hoc multiple comparison test. Effect sizes (η^2^) are reported where appropriate. Data are presented as mean ± SEM. Statistical significance was set at p < 0.05. All behavioral, histological, and molecular analyses were performed by investigators blinded to group assignments.

## Results

### Ultrasound Stimulation Enhances Recognition Memory

To assess the cognitive effects of LIFU stimulation, rats underwent the NOR test before and after the four-week ultrasound intervention. Quantitative analysis revealed that all LIFU-treated groups spent significantly less time in the familiar zone compared to controls (Fig. 2A), with the most pronounced reduction observed in the 0.5 MHz group (*p* < 0.0001), followed by significant decreases at 1 MHz and 5 MHz (*p* < 0.0001). Notably, only the 0.5 MHz group demonstrated a significant increase in time spent in the novel zone (p < 0.0001, Fig. 2B), whereas the 1 MHz and 5 MHz groups showed a non-significant increase. Analysis of novel object-directed behaviors further supported this pattern. Orientation time toward the novel object, defined as the amount of time the animal spent directing its head or snout toward the object while remaining at a distance, was significantly elevated in the 0.5 MHz group (p < 0.0001, Fig. 2C), with moderate but still significant increases observed in the 1 MHz and 5 MHz groups (p < 0.01, Fig. 2C). Interestingly, the time spent getting closer to the novel object, measured as the time the animal physically moved toward or entered the immediate vicinity of the object, was significantly reduced in both the 0.5 MHz (*p* < 0.0001) and 1 MHz (*p* < 0.01) groups, and showed a mild but non-significant reduction at 5 MHz compared to Sham controls (Fig. 2D).

**Fig. 2.**
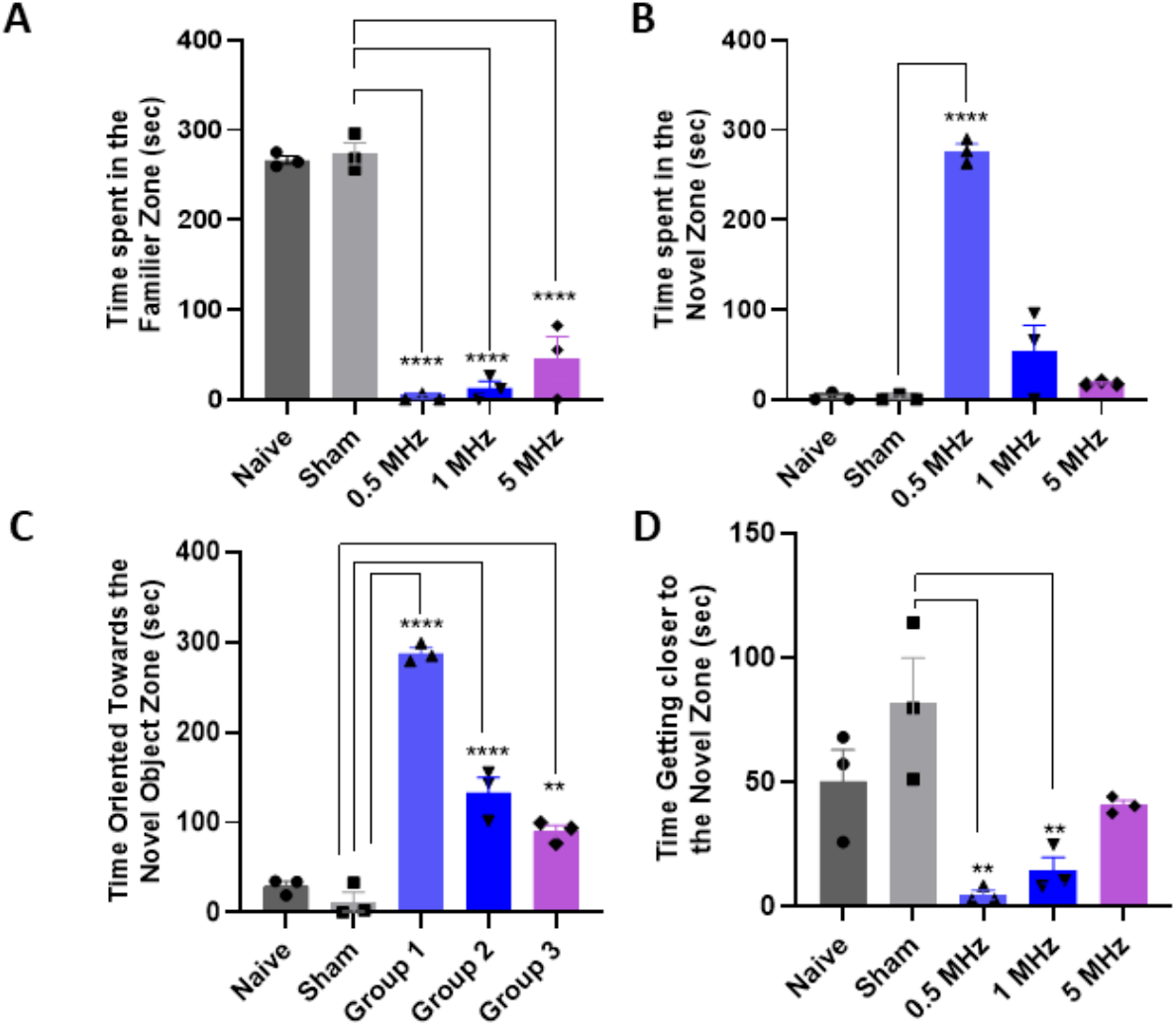
Performance of the rats in the NOR test. (A–D) Quantification of behavioral outcomes, including time spent exploring the familiar zone (A), time spent in the novel object zone (B), time oriented toward the novel object zone (C), and time spent in close proximity to the novel object (D). Across all behavioral measures, ultrasound-treated groups demonstrated enhanced recognition memory compared to Sham, with the 0.5 MHz group showing the most pronounced effects. Statistical significance between assessed using One-way ANOVA followed by post hoc multiple comparison test (**p<0.01,****p<0.0001). Data are presented as mean ± SEM.

### Ultrasound Enhances Neural Stem Cell Proliferation

Across all hippocampal regions, 0.5 MHz LIFU produced the greatest increase in cell proliferation relative to all other groups (Fig. 3A). In the rostral dentate gyrus, the number of BrdU-labeled cells in the 0.5 MHz group (4,296 ± 276, p < 0.001) was significantly greater than that in the sham group (772 ± 77). A similar pattern was observed in the intermediate dentate gyrus, where 0.5 MHz stimulation (1,794 ± 130, p < 0.001) yielded significantly higher counts than all other groups. In the caudal region, 0.5 MHz sonication (1,332 ± 112, p < 0.001) likewise produced significantly greater proliferation compared to the sham group. These regional effects are consistent with the cumulative data shown in Fig. 3A, where the total count of BrdU-labeled cells across all dentate gyrus regions was significantly higher in the 0.5 MHz group (7,422 ± 425, p < 0.0001) compared to the sham (1,624 ± 226) and all other treatment groups. Representative confocal images further confirmed this effect (Fig. 3B), with the 0.5 MHz group showing a clear increase in BrdU^+^/NeuN^+^ co-labeled cells within the DG relative to all other conditions, highlighting that proliferating cells were integrating into the neuronal population.

**Fig. 3.**
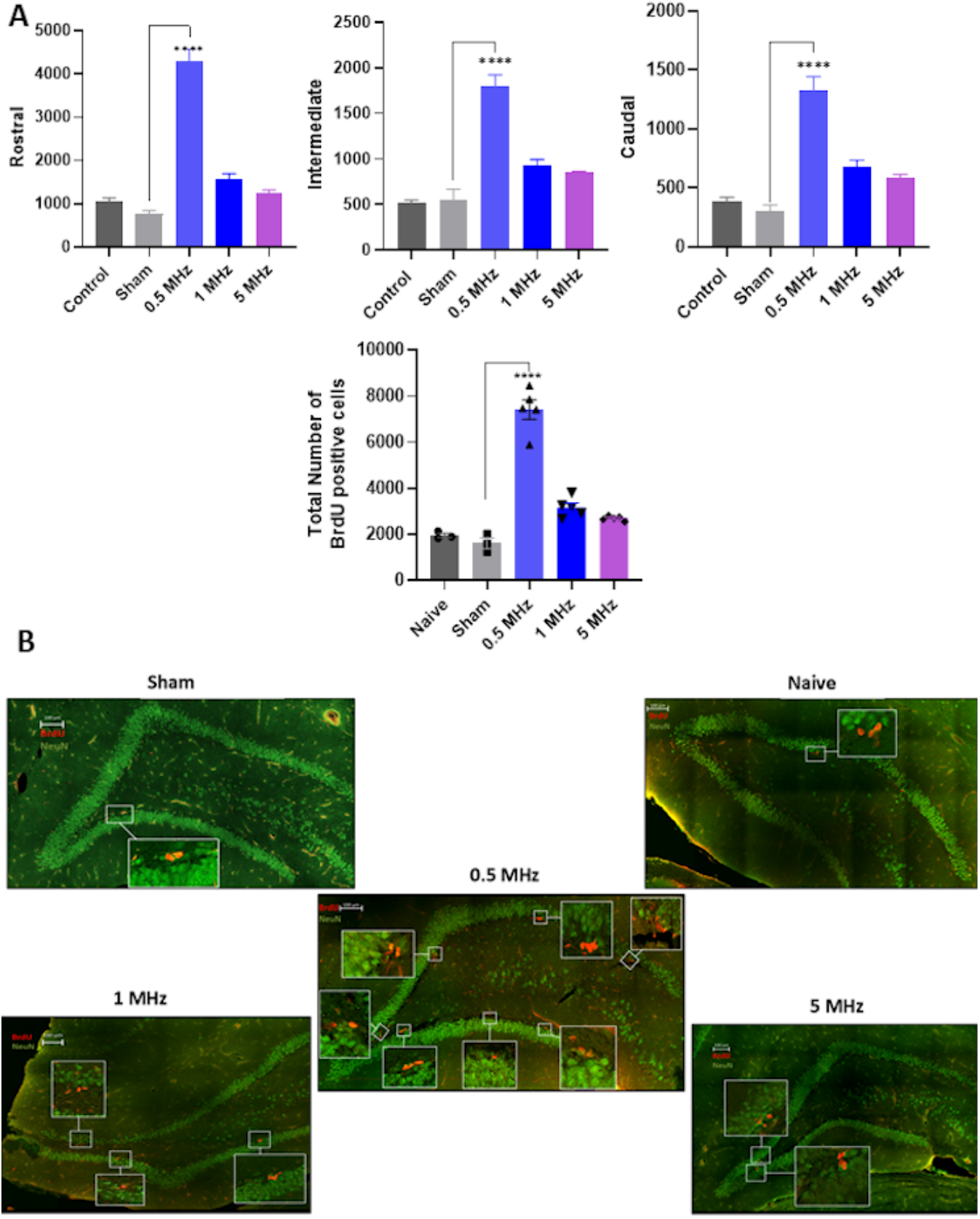
Frequency-dependent increase in BrdU^+^ cell proliferation in the dentate gyrus following low-intensity focused ultrasound stimulation. (A) Stereological quantification of BrdU-labeled cells in the rostral, intermediate, and caudal segments of the DG across experimental groups. The final graph shows the total number of BrdU^+^ cells quantified across the entire DG. Each bar represents the mean ± SEM, and each dot represents the measured value from an individual animal. (B) Representative confocal images showing immunofluorescence labeling of BrdU^+^ cells (red) and NeuN^+^ neurons (green) in the DG from each group. Images were acquired as Z-stacks using a 40× oil objective. A significant increase in BrdU^+^ proliferating cells was observed in the 0.5 MHz group compared with all other groups. Statistical significance was assessed using one-way ANOVA followed by post hoc multiple-comparison testing. ****p < 0.0001

### Ultrasound Stimulation Upregulates Neurogenic Markers

To examine molecular changes associated with ultrasound-induced neurogenesis, we quantified hippocampal mRNA expression of three key neurogenic regulators: BDNF, FGF-2, and Sox-2. Fold-change analysis relative to the Sham group revealed a significant upregulation of all three genes in the 0.5 MHz ultrasound-treated group. Specifically, BDNF expression was significantly increased in the hippocampi of rats treated with 0.5 MHz ultrasound (3.91 ± 0.07, p < 0.0001) compared to Sham (1.02 ± 0.13), whereas the 1 MHz (2.15 ± 0.43) and 5 MHz (1.87 ± 0.36) groups exhibited modest, non-significant increases (Fig. 4A). A similar trend was observed for FGF-2 mRNA, which was significantly upregulated at 0.5 MHz (4.48 ± 0.16, p < 0.0001) relative to Sham (1.12 ± 0.21), while the 1 MHz (2.02 ± 0.39) and 5 MHz (1.74 ± 0.25) groups showed non-significant elevations (Fig. 4B). Sox-2 expression followed the same pattern, showing a significant increase in the 0.5 MHz group (4.21 ± 0.12, p < 0.0001) compared to Sham (1.15 ± 0.19), with only modest, non-significant increases in the 1 MHz (2.11 ± 0.36) and 5 MHz (1.93 ± 0.33) groups (Fig. 4C).

**Fig. 4.**
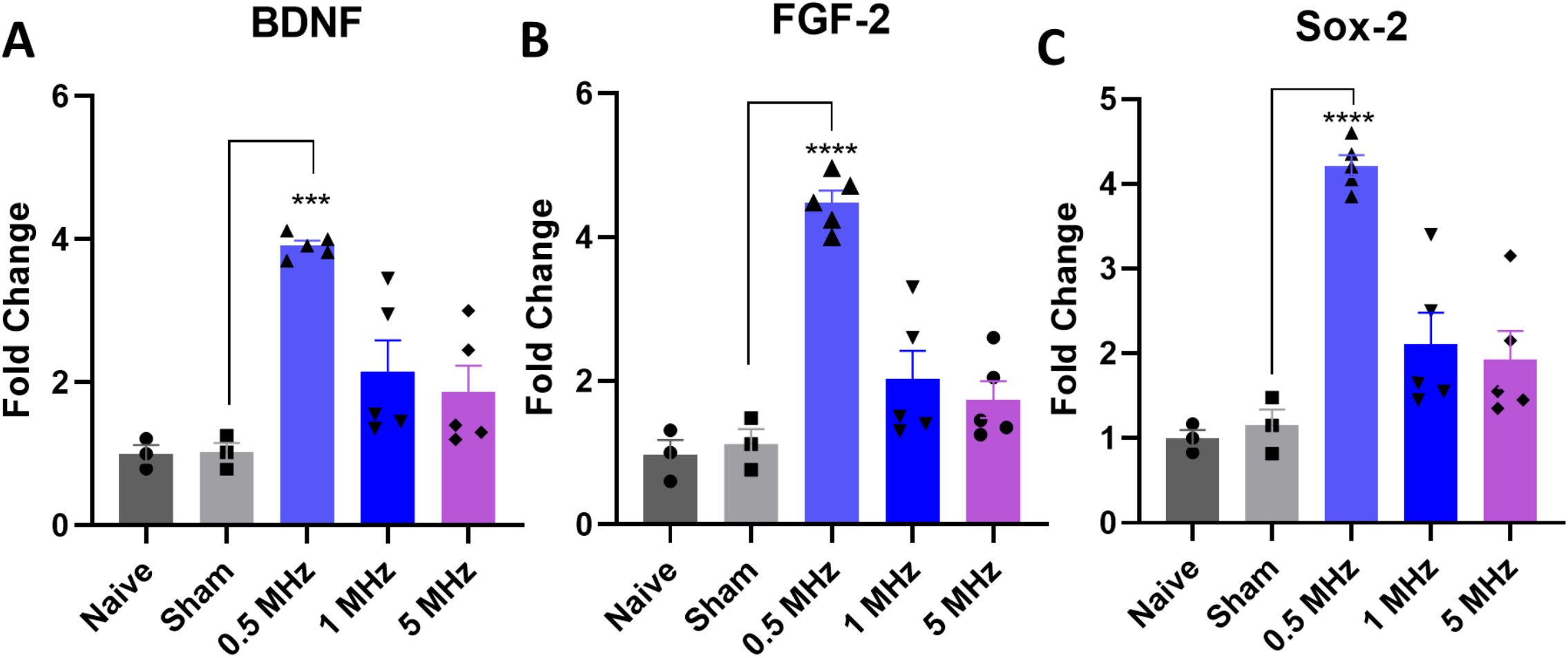
Frequency-dependent effects of LIFU on gene expression. Quantitative PCR analysis revealed that LIFU at 0.5 MHz significantly increased mRNA expression of (A) BDNF, (B) FGF-2, and (C) Sox-2 in the hippocampus. Statistical significance was assessed using one-way ANOVA followed by post hoc multiple comparison testing (***p < 0.001). Data are presented as mean ± SEM.

## Discussion

In the present study, we demonstrate that transcranial low-intensity focused ultrasound (LIFU) enhances adult hippocampal neurogenesis in a frequency-dependent manner. Among the tested parameters, 0.5 MHz stimulation produced the most robust increase in BrdU-labeled proliferating cells across the dentate gyrus, accompanied by elevated expression of BDNF, FGF-2, and Sox-2, and improved recognition memory. These findings indicate that stimulation frequency critically determines the magnitude of ultrasound-induced hippocampal plasticity.

A central observation of this study is that lower-frequency stimulation resulted in greater neurogenic and molecular responses despite identical spatial-peak pulse-average intensity across groups. Frequency influences the mechanical characteristics of tissue displacement under acoustic pressure. Lower frequencies generate larger oscillatory displacements, which may increase membrane deformation and engagement of mechanosensitive signaling pathways. In neural progenitor populations, mechanotransduction has been shown to regulate proliferation and differentiation through calcium-dependent signalling cascades, including activation of Piezo1, TRP channels, and voltage-gated calcium channels. Downstream activation of MAPK/ERK and CREB pathways promotes transcription of neurotrophic factors such as BDNF, which are essential for adult hippocampal neurogenesis. The observed upregulation of BDNF, FGF-2, and Sox-2 in the 0.5 MHz group is consistent with involvement of these pathways.

Although the ultrasound field cover the dentate gyrus along its rostrocaudal extent, the proliferative response was not uniform across regions. In our sectioning scheme, the rostral dentate gyrus corresponds predominantly to the dorsal hippocampus, whereas the intermediate and caudal regions extend toward more ventral hippocampal domains. The stronger proliferative response observed in the rostral region is therefore broadly consistent with the improvement in recognition memory detected in the NOR task, given the established role of dorsal hippocampal circuits in learning and memory.

The mechanical strain induced by low-frequency ultrasound may engage calcium-permeable channels such as Piezo1, members of the transient receptor potential (TRP) family [40, 41], and voltage-gated calcium channels (VGCCs)[42]. Such mechanical stimulation has been proposed to influence intracellular signaling cascades including MAPK/ERK [43, 44], PI3K/Akt [45], NF-κB [46], and CREB pathways, although these pathways were not directly examined in the present study. Previous in vitro work has demonstrated that transient ultrasound exposure can induce dose-dependent phosphorylation of ERK1/2 (Thr202/Tyr204), partially mediated by oxidative signalling mechanisms, indicating that ultrasound can engage intracellular signalling cascades under specific conditions [47]. In addition, alterations in BDNF/TrkB signalling have been linked to hippocampal function in vivo, supporting the biological relevance of this pathway [48]. These pathways may influence transcription factors that regulate proliferation and survival, ultimately driving expression of neurotrophic and stemness-associated genes [49]. The upregulation of BDNF, FGF-2, and Sox-2 observed in the present study is therefore compatible with activation of neurogenic signalling; however, these findings remain correlative, and no direct causal pathway can be inferred. Importantly, Sox-2 expression points to sustained maintenance of the neural stem cell pool, suggesting that repeated ultrasound not only promotes proliferation but also preserves long-term neurogenic potential [50]. Mechanical stress can directly alter neuronal function by deforming the cell membrane and cytoskeleton, thereby modulating mechanosensitive ion channels, disrupting ion homeostasis, and triggering biochemical signaling pathways that can influence excitability, synaptic transmission, survival, or injury responses[51, 52].

The molecular and cellular enhancements induced by low-frequency ultrasound were accompanied by improved recognition memory. In the NOR task, 0.5 MHz stimulation increased novelty exploration and orientation toward new objects, reflecting hippocampal-dependent discrimination. The fact that these cognitive benefits developed after multiple weeks of stimulation suggests that cumulative processes including neural progenitor proliferation, differentiation, and integration are required for behavioral expression. These processes are likely reinforced by sustained upregulation of neurotrophic and neuroplasticity-related genes (BDNF, FGF-2, Sox-2), which promote neurogenesis, synaptic remodeling, and long-term potentiation (LTP), all critical for memory formation [15, 53]. This aligns with prior studies showing that multi-session ultrasound protocols are necessary to induce durable plasticity and behavioral gains [44].

Our findings build on prior studies that ultrasound can upregulate neurotrophic factors and improve cognition in models of aging and neurodegeneration. Early clinical studies have similarly demonstrated cognitive improvements in AD patients following hippocampal ultrasound, even without blood–brain barrier opening [54, 55]. In parallel, preclinical work by Shimokawa et al. [56] and Nicodemus et al. [57] reported sustained cognitive improvements following multi-session ultrasound stimulation, further supporting the relevance of repeated sonication for future preclinical evaluation. By identifying frequency as a key driver of efficacy, our results provide a framework for optimizing ultrasound protocols to maximize neurogenic and cognitive outcomes.

Nonetheless, several limitations should be acknowledged. First, the molecular findings are correlative, as increased BDNF, FGF-2, and Sox-2 mRNA expression was not accompanied by protein-level validation or assessment of downstream signaling pathways; thus, functional engagement of neurogenic signaling cascades cannot be confirmed. Second, given that BrdU labeling was assessed 24 h after administration, the cellular data mainly reflect recent proliferative activity and do not fully capture later stages of neurogenesis such as immature neuronal development, survival, or maturation; additional markers such as Ki-67 and DCX would help further characterize the neurogenic processes influenced by LIFU. Third, the study design may limit generalizability, as only male rats were included. Adult hippocampal neurogenesis and BDNF signaling are known to be modulated in a sex-dependent manner, with ovarian hormone fluctuations influencing progenitor cell proliferation, neuronal differentiation, and neurotrophic pathways [58]. Therefore, the present findings may not fully generalize to females. Add to that, sample sizes were modest, particularly in the control groups, and experiments were conducted in healthy adult animals rather than in a disease model. Fourth, we acknowledge that behavioral assessment was limited to the NOR test, which, although widely used and sensitive to hippocampal dysfunction under specific conditions, does not fully capture the range of hippocampal-dependent cognitive processes. Future studies should incorporate additional tasks, such as the Morris Water Maze or Contextual Fear Conditioning, to provide a more comprehensive assessment of spatial and associative memory and to further validate the functional relevance of LIFU-induced neurogenic effects, especially given that the outcomes were measured immediately after the 4-week stimulation period, so broader functional effects and the durability of the observed changes remain to be established.

Regarding our acoustic simulation approach, simulations were performed using a simplified 2D water-skull model, which primarily captures skull-induced attenuation and phase distortion. While this approach is appropriate for comparing relative frequency-dependent transmission across transducers, it does not account for three-dimensional skull geometry or intracerebral tissue heterogeneity. Our group previously validated the accuracy of this simulation approach through experimental acoustic characterization, including hydrophone measurements, demonstrating agreement with simulated results with a deviation of less than 10% [34]. Nevertheless, future studies should incorporate 3D modeling frameworks and anatomically realistic skull representations to improve spatial accuracy and translational relevance.

In summary, these data demonstrate that ultrasound frequency is a critical determinant of adult hippocampal neurogenic and cognitive responses. By identifying frequency-dependent modulation of neurotrophic signaling and neural progenitor proliferation, this study provides a biological framework for investigating ultrasound-mediated regulation of hippocampal plasticity under physiological and pathological conditions.

## Conclusion

In summary, low-intensity focused ultrasound enhances adult hippocampal neurogenesis and recognition memory in a frequency-dependent manner, with lower frequencies producing the most pronounced cellular and molecular effects. These findings identify stimulation frequency as a critical determinant of ultrasound-induced modulation of hippocampal plasticity. However, as the study was conducted in healthy animals, the relevance of these effects to pathological states, including neurodegenerative disorders, remains to be established. Therefore, the current results should be interpreted as foundational, and future studies targeting disease-relevant models are required to determine the therapeutic potential and translational applicability of LIFU-mediated neuromodulation.

## Data Availability

Data used in this study are available from the corresponding author upon reasonable request.

## Author Contributions

**Kareen Kanaan:** Conceptualization, Methodology, Investigation (animal experiments, histology), Writing – Original Draft (co-wrote with H. Badawe).

**Heba Badawe:** Methodology, Investigation (animal experiments), Writing – Original Draft (co-wrote with K. Kanaan).

**Wassim Abou-Kheir*:** Resources, Supervision, Writing – Review & Editing.

**Massoud L. Khraiche*:** Conceptualization, Supervision, Funding Acquisition, Writing, Review & Editing,

## Authors’ contributions

- **Kareen Kenaan:** Performed the experiments, collected the data, carried out all analyses, and contributed to writing the manuscript.
- **Heba Badawe:** Performed experiments, collected part of the data, carried out partial analyses, and contributed to writing the manuscript.
- **Wassim Abou-Kheir**: Supervised the project and provided critical review and editing of the manuscript.
- **Massoud Khraiche:** Supervised the project, contributed to the study design, and assisted in writing and editing the manuscript.

## Consent for publication

Not applicable. This study does not include individual human data or identifiable images requiring consent.

## Ethics approval and consent to participate

All protocols were approved by the Institutional Animal Care and Use Committee (IACUC) of the American University of Beirut. Protocol # 237808.

## Acknowledgments

Not applicable.

